# Enhancement of compounds in hydrosol and residual water of *Aquilaria malaccensis* tetraploid

**DOI:** 10.1101/492405

**Authors:** A.R. Siti Suhaila, N. Mohd Saleh, M. Norwati, P. Namasivayam, C. Mahani, K.I. Kandasamy

## Abstract

*Aquilaria malaccensis* is an agarwood-producing species in the family Thymeleaeceae. Agarwood is a fragrant resin used in the manufacture of incense sticks, and in pharmaceutical, perfumery and cosmetic industries. In addition to the resin, hydrosol and residual water by-products from agarwood woodchip distillation are also utilized. Hydrosol contains water-soluble fragrant chemicals used as a tonic drink, in cooking and cosmetics while the residual water is used in spas and aromatic bath treatments. The present study was conducted to identify and compare compounds present in hydrosol and residual water by-products of diploid and polyploid *A. malaccensis*. Four different four-month-old *A. malaccensis* plants were compared: soil-grown diploid seedlings (DS), *in vitro*-grown seedlings (DV), tissue culture-derived plantlets (DC) and artificially induced tetraploid plantlets (TC). Hydrosol water from TC leaf and root samples were found to contain higher amounts of compounds compared with other samples. The TC leaf samples were qualitatively better as key compounds of agarwood such as α-and γ-eudesmol were detected. TC stem samples also contained higher amounts of key compounds compared with other samples, while the overall amount of compounds was highest in DS stem samples. The residual water of TC stem and root samples contained key compounds not detected in other samples, while DS residual water samples contained the highest total amount of compounds. *Aquilaria malaccensis* tetraploids performed better than their diploid counterparts in production of compounds, and thus may be a better planting material choice for commercial plantations.

## INTRODUCTION

*Aquilaria malaccensis* (Thymealaeceae) is one of 25 *Aquilaria* species (agarwood-producing species) distributed throughout Peninsular Malaysia, Indonesia, India, Bangladesh and Thailand [15, 16]. Their agarwood has been traded across Europe, the Middle East and Asia for more than 2000 years. It is considered to be one of the most expensive natural products existing today and is known among agarwood traders as the ‘black gold of the forest’ used to make valuable products such as pure agarwood oil, cosmetics, and medicines and incense [15]. How agarwood is formed is not yet fully understood, but the general understanding is that it is formed and permeates the heartwood as a stress response to wounding, microbial infection or both [15, 21]. Agarwood yield increases as the trees get older, and infected trees are able to produce high quality agarwood oil at the age of 20 years [4]. The price of agarwood oil increased tremendously from MYR280/tola (12 ml) at the end of 1990s to MYR500/tola recently. AAAAA grade agarwood was recently priced between MYR120,000–360,000/kg. While these high prices benefit traders, the exponential demand for agarwood and illegal harvesting of wild *A. malaccensis* trees from natural forest parks and reserves has caused this species to be listed under CITES (the Convention on International Trade in Endangered Species of Wild Fauna and Flora, Appendix II) [21].

Sesquiterpenoids and phenylethyl chromone derivatives are the principal compounds found in the agarwood oil, with α-guaiene, δ-guaiene and α-humulene making up the major volatile compounds [15, 16]. In addition to agarwood essential oil, the water-soluble fraction called hydrosol (also known as floral water) is a valuable by-product obtained by hydro distillation. Hydrosol contains all the agarwood essential oils in low concentrations, making it suitable for making products where pure essential oil would be considered too strong. Agarwood-based products such as soap, perfume water and drinking water usually incorporate hydrosol water and residual water. As wild *A. malaccensis* populations in natural forests shrink and agarwood demand increases, non-conventional methods in agarwood production need to be explored. Chromosome doubling is known to have an effect on many physiological properties of a plant, e.g. producing larger reproductive and vegetative plant parts [5, 6, 7, 10, 11] and enhancing secondary metabolite production [7, 10]. Plants such as *Artemisia annua* L., Vetiver and Papaver species exhibit increased key secondary metabolite content [6, 8, 9]. Artemisinin content enhancement in *A. annua* was achieved by polyploidization in its hairy roots. *Datura* and *Papaver* plant species showed very high (almost 100%) increases in ssecondary metabolites after chromosome doubling [6, 10]. Crops are also known to benefit from the hydrosol water and residual water, with improvements seen in the yield and quality of some essential oil crops [24]. Enhancing the compounds in young *A. malaccensis* may help reduce illegal harvesting of the species. The present study was conducted to identify and compare compounds present in hydrosol and residual water by-products of diploid and polyploid *A. malaccensis*.

## MATERIALS AND METHODS

### Plant materials

Four different four-month-old *A. malaccensis* plant materials were compared: soil-grown diploid seedlings (DS), *in vitro*-grown seedlings (DV), *in vitro*-grown tissue culture-derived plantlets (DC) and *in vitro*-grown artificially-induced tetraploid plantlets, polyploids (TC). Plant parts (leaf, stem and roots) were separated and subjected to hydro distillation processes.

### Hydro distillation process

Plant parts (leaf, stem and root) from all plant samples were weighed at 20 g and placed into individual round-bottomed flasks containing 0.5 L of distilled water. The samples were boiled for 6 hours at 100 °C. The distilled agarwood compounds were collected in a Clevenger column. Hydrosol water (water condensed and trapped together with agarwood oil in the Clevenger column) and residual water (water that was left in the flask) was evaluated to determine the compounds present.

### Gas Chromatography-Mass Spectrometry

Samples collected in the Clevenger column were heated at 60 °C for 10 minutes at a rate of 3 °C per minute. The temperature was gradually increased by 68–77 °C to 230 °C and held at that temperature for 10 minutes. Gases that were released from samples were then injected into a gas chromatography-mass spectrometry (GC-MS) instrument. Different compounds vaporized at different times (‘retention time’, RT). GC-MS was used to analyse compounds detected at their respective peaks. The mass spectrometer peaks that were considered for analyses were those that at least more than 90 percent matched to the library. The peaks were identified by comparing with mass spectrometer of the HPCH 2205.L, Wiley7 NiST05. And NIST0.5a.L. Retention times are used to calculate relative retention times—dividing the retention time of a compound by the retention time of an internal standard. Tables of retention indices (RI) were used by comparing experimentally found retention indices with known values. The RI is given by the equation:

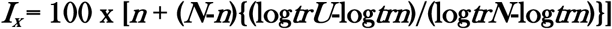

***x*** : the name of the target compound
*n0* : n-alkane Cn0H2n0+2 directly eluting before ***x***
***n1*** : n-alkane Cn1H2n1+2 directly eluting after ***x***
***RT*** : retention time (in any unit such as minutes, seconds, etc.)
***RI*** : retention index (pure number without unit)

### Statistical Analyses

Mean percentage values (± SD) were used to determine the compounds present in the samples using solid-phase microextraction (SPME)-GCMS. Peaks were identified by comparing the retention times of the samples with those from the National Institute of Standards and Technology database.

## RESULTS AND DISCUSSION

### Analyses of hydrosol

Hydrosol from TC leaves showed the highest peak area for compounds with more than 10 fold higher than that from DC, DS, and DV (Table 1). In TC leaf samples, five important compounds were detected, i.e. α-, γ-and 10-epi-γ-eudesmol, epi-α-muurolol and δ-selinene. Eudesmol are found in high quality agarwood essential oil (15, 16). Several important compounds were detected in TC leaf samples, but in smaller amounts, i.e. α-and γ-muurolene, α-selinene and γ-gurjunene. Two important compounds were detected in DS leaf samples, i.e. α-and γ-muurolene. Chromosome doubling in *TC* leaves may have resulted in the high amounts of compounds detected in the hydrosol from those samples.

**Table 1.** Compounds in hydrosol water of *A. malaccensis* from field-grown seedling, *in vitro* seedlings, *and in vitro* diploid and tetraploid plantlets

| Compound | RI | Leaf samples: |  |  |  | Stem samples: |  |  |  | Root samples: |  |  |  |
| --- | --- | --- | --- | --- | --- | --- | --- | --- | --- | --- | --- | --- | --- |
|  |  | DS | DV | DC | TC | DS | DV | DC | TC | DS | DV | DC | TC |
| allo-Aromadendrene | 1517 | - | - | - | - | 1.91±3.30 | - | - | 0.54±0.94 | - | - | - | - |
| Aromadendrene | 1599 | - | - | - | - | - | - | - | 2.40±4.16 | - | - | - | - |
| Bicyclogermacrene | 1494 | - | - | - | - | - | - | - | - | - | - | - | 3.20±0.74 |
| α-Cadinol | 1632 | - | - | - | - | - | - | - | - | - | 19.25±0.78 | - | - |
| *trans*-Cadin-1(6),4-diene | 1475 | 1.23±2.14 | - | - | - | - | - | - | - | - | - | - | 0.47±0.98 |
| γ-Cadinene | 1513 | 0.84±1.46 | - | 0.77±1.34 | - | - | - | - | 0.12±0.21 | - | - | - | - |
| β-Calacorene | 1564 | - | - | - | - | 3.97±3.84 | - | - | - | - | - | - | - |
| *cis*-Calamenene | 1590 | - | - | - | - | 2.50±2.90 | 0.98±2.23 | - | - | - | 0.31±0.01 | - | - |
| *trans*-Calamenene | 1592 | - | - | 0.66±1.15 | - | - | - | - | - | - | - | - | - |
| (E)-Caryophyllene | 1408 | - | - | - | - | - | - | - | - | - | - | - | 1.31±0.20 |
| 9-*epi*-(E)-Caryophyllene | 1536 | - | - | - | - | - | - | - | 1.14±1.97 | - | - | - | - |
| 1,8-Cineole | 1011 | - | - | - | - | - | - | - | - | - | 11.81±3.88 | - | - |
| Citronellyl Acetate | 1346 | - | - | - | - | - | - | - | - | - | - | - | 1.41±1.20 |
| Cuparene | 1502 | - | - | - | - | - | - | - | - | - | - | - | 0.32±5.54 |
| *ar*-Curcumene | 1479 | - | - | - | - | - | - | - | 0.67±1.17 | - | - | - | - |
| Curazene | 1493 | - | - | - | - | - | - | - | - | - | - | - | 1.09±1.02 |
| Elemol | 1538 | - | - | - | - | - | - | 1.24±2.14 | - | - | 3.93±0.43 | - | - |
| 10-*epi*-γ-Eudesmol | 1563 | - | - | - | 0.58±1.00 | - | - | - | - | - | - | - | - |
| α-Eudesmol | 1635 | - | - | - | 12.34±21.4 | - | - | - | - | - | - | - | - |
| γ-Eudesmol | 1632 | - | - | - | 13.97±6.01 | - | - | 1.17±2.02 | - | - | 6.11±1.98 | - | - |
| Geranyl Butanoate | 1377 | - | - | - | - | - | - | - | - | - | - | - | 5.96±2.84 |
| Germacrene D | 1492 | - | - | - | - | - | - | - | - | - | - | - | 0.44±0.98 |
| α-Guaiane | 1535 | - | - | - | - | - | - | - | 0.63±1.08 | - | 0.27±2.76 | - | - |
| β-Gurjunene | 1542 | - | - | - | - | - | - | - | 1.03±0.95 | - | - | - | - |
| γ-Gurjunene | 1530 | - | - | 0.29±0.50 | - | - | - | 1.66±2.88 | 0.32±0.89 | - | - | - | - |
| α-Gurjunene | 1509 | - | - | - | - | - | - | 9.33±12.12 | 0.72±1.24 | - | - | - | - |
| Hedycaryl | 1546 | - | - | - | 15.39±6.41 | - | - | 0.45±0.78 | - | - | - | - | - |
| E-β-Ionone | 1487 | - | 0.40±1.20 | - | - | - | - | - | - | - | - | - | - |
| Longifolene | 1679 | - | - | - | - | - | - | - | 5.50±5.01 | - | - | - | - |
| Methyl dodecanoate | 1594 | - | - | - | 0.16±0.27 | - | - | - | - | - | - | - | - |
| Methyl γ-ionone | 1479 | - | - | - | - | - | - | - | 0.31±0.40 | - | - | - | - |
| *epi*-α-Muurolol | 1572 | - | - | - | 3.71±6.42 | - | - | - | - | - | 7.55±2.04 | - | - |
| α-Muurolene | 1441 | 0.69±1.20 | - | 1.13±1.96 | - | - | - | - | - | - | - | - | - |
| γ-Muurolene | 1425 | 0.33±0.56 | - | 0.79±1.37 | - | - | - | - | - | - | 0.38±1.09 | - | 3.45±0.32 |
| Pentadecane | 1500 | - | - | - | - | - | - | - | 0.47±0.82 | - | - | - | - |
| Prenanaspodiene | 1669 | - | - | - | - | - | - | - | 0.63±0.18 | - | - | - | - |
| Rosifolol | 1600 | - | - | - | - | - | - | 0.91±0.58 | - | - | - | - | - |
| Selin-11-en-4-α-ol | 1667 | - | - | - | - | - | - | 3.35±5.88 | - | - | - | - | - |

DS=field-grown seedlings, DV=*in vitro* seedlings, DC=*in vitro* diploid, TC=tetraploid plantlets, RI=retention index on the HP-5MS capillary column of a homologous series of $C_6$ - $C_{18}$ *n*-alkanes; - : absent.

Among stem samples, DS had the highest peak area of compounds followed by DC, TC and DV (Table 1). However, DS stems contained a very high level of detectable compounds, only 1.91% of these were important compounds while the rest were aliphatic hydrocarbons except for allo-aromadendrene. On the other hand, DC contained highest level of important compounds followed by TC. Compounds such as α-and γ-gurjunene were detected in DC stems and in lower amounts in TC stems. Other important compounds detected in TC stems were γ-muurolene, α-and δ-selinene, and cis-β-guaiene.

The roots of all samples showed the presence of γ-gurjunene with the highest level found in TC followed by DS, DV and TC (Table 1). Several important compounds were detected in DS roots, i.e. *cis*-β-guaiene and α-agarofuran, while DC contained low levels of α-muurolene and aromadendrene compounds detected at a high level in DC was eugenol, which although not an important sesquiterpene in agarwood oil, is valuable as an antiseptic and anesthetic compound in the pharmaceutical industry. Eugenol can be combined with zinc oxide to form zinc oxide eugenol, which has restorative and prosthodontic applications in dentistry. TC contained low levels of other important compounds, i.e. α-and γ-muurolene, α-and δ-selinene, and cis-β-guaiene.

### Analyses of residual water

The residual water of only DV and TC contained compounds, with higher levels found in compounds from TC (Table 2). DV leaf samples had the highest total peak area and the important compounds α-guaiene and γ-eudesmol were present. TC leaves contained a lower level of detectable compounds with a sole important chemical constituent, γ-muurolene. Among the stem samples, a total peak area of 5.90% was detected in DV and 1.83% in TC. γ-Eudesmol is an important chemical constituent that was detected in DV. Again, no sesquiterpenes were detected in DS and TC. Among the root tissue samples, DV contained a higher total peak area than TC compounds but almost 50% of the total compounds detected in TC were important, compounds i.e. α-muurolene (0.91%) and δ-selinene (Table 2). DV did not have any important compounds, and no compounds were detected in DS and TC residual water root samples. The low or undetectable amounts of compounds in residual water indicate most of the chemicals were released during hydro distillation. It is notable that a few important compounds were found in DV and TC residual water samples compounds after hydro distillation.

The thicker cell walls of DC compared with those of TC may have resulted in retention of these chemical constituents during hydro distillation. In line with the higher levels of compounds found in TC as compared with diploid plants in the present study, previous research has also shown that polyploidization can enhance secondary metabolite production in plants [e.g. 6, 4 and 10]. Another major factor influencing secondary metabolite production is growth condition. Seedlings grown under natural conditions contained the high level of total volatile compounds compared to plant tissues from those grown under control conditions (*in vitro*) [1, 20]. Production of secondary metabolites can also increase in the presence of higher light intensity whereby many such compounds are important in plant defense and produced in response to external threat stimuli [12, 13]. Therefore lower levels may be produced in *in vitro* plants that are unlikely to be attacked or damaged by biotic and/or abiotic factors.

The *in vitro* tetraploids in our study produced high levels of important sesquiterpenes, indicating higher gene expression levels in the tetraploid than in the diploid plants. The large quantities of important compounds detected in the hydrosol from all plant samples showed that hydro distillation can efficiently extract compounds from *A. malaccensis* compounds. The higher yields of secondary metabolites demonstrated by the tetraploid *A. malaccensis* in our study are consistent with those reported for the tetraploid rose (*Rosa damascene*), which is widely used to produce rose oil and rose water in quantity, and is probably triparental in origin [17,23]. Our results indicate that while the total quantity of compounds was about the same in diploid and tetraploid *A. malaccensis*, the composition of these compounds was of better quality in tetraploids, which had relatively higher levels of important sesquiterpenes. Cells with higher ploidy levels have larger genome sizes, and are expected to have higher levels of transcription than cells with lower ploidy levels [2, 5, 10]. Therefore, gross expression of most or some of these genes would be expected to increase linearly with ploidy level. Previous studies have demonstrated that different parts of plants produce different secondary metabolites [11, 19, 20, 22] and the results of our study are in line with this different compounds were detected in the leaves, stems and roots of diploid and tetraploid *A. malaccensis*. The enhanced production of important compounds in *in vitro* tetraploids can potentially add more value to *A. malaccensis* than its diploid counterparts. Commercial planting of tetraploid *A. malaccensis* yielding higher amounts of these important compounds than the current diploid plantings.

## Acknowledgments

The authors thank FRIM for financial support through RMK10 (Vot#10310802005) and Universiti Kebangsaan Malaysia for use of experimental facilities (flow cytometer laboratory).

